# Evolutionary conservation of mechanical strain distributions in functional transitions of protein structures

**DOI:** 10.1101/2023.02.07.527482

**Authors:** Pablo Sartori, Stanislas Leibler

## Abstract

One of the tenets of molecular biology is that dynamic transitions between three-dimensional structures determine the function of proteins. Therefore, it seems only natural that evolutionary analysis of proteins, presently based mainly on their primary sequence, needs to shift its focus towards their function as assessed by corresponding structural transitions. This can be facilitated by recent progress in cryogenic electron microscopy that provides atomic structures of multiple conformational states for proteins and protein assemblies isolated from evolutionarily related species. In this work, we study evolutionary conservation of multi-protein assembly function by using mechanical strain as a quantitative footprint of structural transitions. We adopt the formalism of finite strain analysis, developed in condensed matter physics, and apply it, as a case study, to a classical multi-protein assembly, the ATP synthase. Our Protein Strain Analysis (PSA) provides a precise characterization of rotation domains that agrees with the present biophysical knowledge. In addition, we obtain a strain distribution on the protein structure associated with functional transitions. By analyzing in detail, the strain patterns of the chains responsible for ATP synthesis across distinct species, we show that they are evolutionarily conserved for the same functional transition. Such conservation is not revealed by displacement or rotation patterns. Furthermore, within each functional transition, we can identify conserved strain patterns for ATP synthases isolated from different organisms. The observed strain conservation across evolutionary distant species indicates that strain should be essential in future structure-based evolutionary studies of protein function.

## Introduction

Evolution acts by natural selection of multifaceted features of living systems, phenotypic traits, that are encoded in their DNA sequence, genotype. At the sub-cellular scale, many phenotypic traits can be associated to the functioning of proteins and multi-protein assemblies [1]. Most approaches for evolutionary comparison of proteins focus on their DNA sequence [2]. While this sheds light into the genotypic space, taking phenotypic traits into account requires comparing proteins as they perform their multiple cellular functions. To study evolution of protein phenotypes, the focus must, therefore, shift towards the evolutionary changes in the different structural transitions underlying protein functions.

Evolutionary comparison of protein structures encounters two fundamental hurdles [3]. First, despite recent efforts [4], spatial alignment remains an arbitrary element of structural comparison. Second, it is not a static structure that defines the function of a protein, but rather dynamical transitions between multiple structural states. Mechanical strain is an alignment-independent quantity that has recently been used to study structural transitions [5, 6], and has therefore the potential to overcome both these barriers. The usage of strain as a probe for protein function is justified because on functional timescales proteins behave largely as elastic materials [7, 8], which explains why elastic paradigms have been so successful in protein biophysics during the last decades [9]. However, despite recent applications to diverse protein structures [10–12], the relationship between strain patterns and rotation or displacement descriptions traditionally associated to protein function remains unaddressed. Here, we establish this relationship, and show that distinct strain patterns correspond to distinct functions performed by one same protein. This allows us to address the fundamental question: are structural transitions of proteins corresponding to different functions evolutionary conserved? In other words, beyond their sequence similarity, can we asses evolutionary conservation of protein function using mechanical strain?

### Protein Strain Analysis of ATP synthase

We adapted the complete formalism of finite strain analysis, which belongs to a long-established branch of physics, elasticity theory [15, 16], to proteins. Our Protein Strain Analysis (PSA, see Appendix A for a detailed method summary) computes local quantities (defined for a neighborhood of each residue taking C_*α*_ as reference) analogous to those of finite strain theory. Examples of such quantities are the principal stretches, which measure the magnitude of strain, or the local rotation angle, which measures structural changes that do not strain the structure. As a case study we chose the ATP synthase. This was motivated by its important functional conservation concomitant with evolutionary variability [17], abundant biophysical and molecular dynamics modelling of its dynamics [18–23], and the recent availability of multiple structural conformations for several ATP synthases isolated from evolutionary distant organisms [24–32]. Furthermore, the well-established connection between ATP synthase function and rotations is an ideal test-ground for PSA.

This protein assembly synthesizes ATP driven by a membrane proton gradient [24, 33, 34] (see Appendix B for detailed schematics). Through its functional cycle the membrane-submerged F_o_ complex, composed of a *c*-ring together with its central “stalk” *γ*, adopts three disctinct rotational states (I, II and III) relative to the rest of the assembly [35, 36]. Proton driven rotation of F_o_ results in interactions with the F_1_ complex, with approximate three-fold symmetry. The F_1_ complex includes three distinct *β* chains *β*_1_, *β*_2_ and *β*_3_, participating in ATP synthesis. In any of the three rotary states, each of the three distinct *β* chains occupies one of three distinct enzymatic (and conformational) states, which we call here E (for the “empty” APO state), D (for “ADP bound”) and T (for “ATP synthesis”). As the whole assembly performs its rotary cycle I *→* II *→* III …, each of the three *β* chains undergoes the enzymatic cycle E *→* D *→* T *→* … [37, 38]. Each of the three transitions of *β* correspond to a different function: ADP binding (E *→* D), ATP synthesis (D *→* T), and ATP release (T *→* E). Therefore, the ATP synthase can intuitively be described as a rotary engine, in which rotation of F_o_ is coupled to conformational transitions of *β*s that perform specific functions.

To provide a quantitative account of this description, we determined the rotation angle and axis for the assembly transition I *→* II (see section B of SI for table of supporting files with information on other transitions, and note that we aligned structures using the channel chain *a*). Fig 1A shows a scatter plot of the local (per residue *i*) rotation angle, *θ*(res_*i*_), against the *z*-component of the rotation vector, *r*_*z*_(res_*i*_). We identify a cluster (cluster 2, in green) that corresponds to a rotation angle *θ ≈* 100°, and maps onto the F_o_ region of the protein, see Fig 1B. This is in agreement with the rotation of 103° reported in [14]. Note that the fact that the rotation angle is not 120° emphasizes the precession and the lack of perfect global symmetry of the structure. In addition, within the chain *β*_3_ that undergoes E *→* D to bind ADP, we also identify two clusters (in orange and purple, Fig 1A), which map into two compact domains of the structure, Fig 1B. These domains are separated at the height of the site of ATP synthesis. Therefore, PSA robustly quantifies local rotations, which are distributed over large compact domains (at multi-chain levels and sub-chain levels). This agrees with the intuitive understanding of how the ATP synthase functions, yet our analysis requires no *a priori* knowledge of the detailed geometrical nature of the structure.

**FIG. 1:**
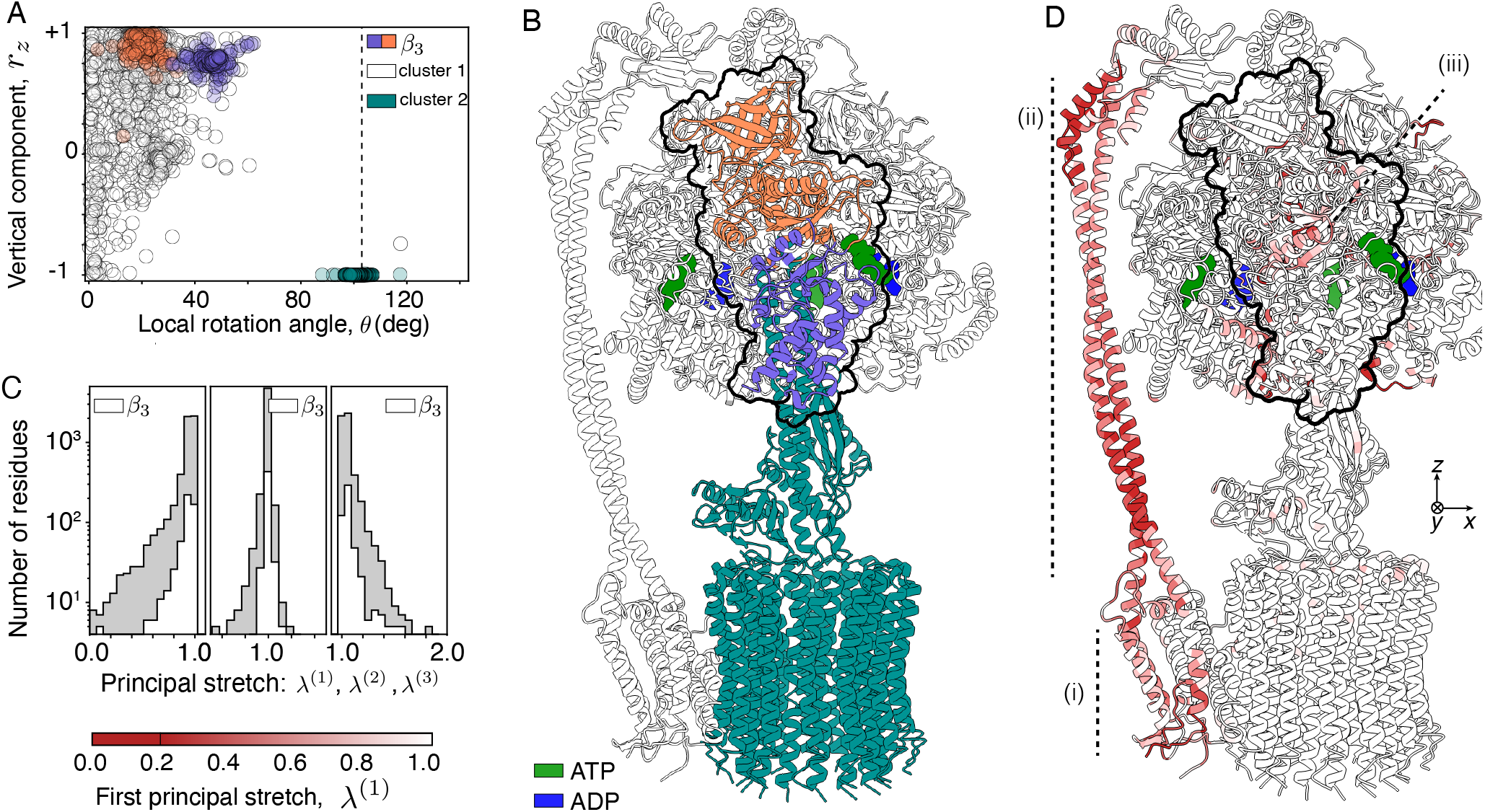
Rotation and strain analysis of ATP synthase structural transition. **A**. Scatter plot of rotation angle *θ* against the vertical component of the rotation axis, *r*_*z*_, for each residue of the structure. Each color labels a cluster obtained using the algorithm in [13] with parameter *ρ* = 0.1. The dark green cluster corresponds to *θ ≈* 100° and agrees with the rotation of 103° reported in [14] for F_o_. Amino acids of *β*_3_ can be separated in two further clusters, one purple and one orange, with distinct rotation angles. **B**. Mapping of the rotation-space clusters on the spatial structure of the synthase. At the whole assembly level, rotations occur in large compact regions corresponding to F_o_ and F_1_. Focusing on *β*_3_, outlined, the two rotation-space clusters map into compact regions of the chain. These regions are separated by an interface at the height where ATP is located. **C**. Histogram of principal stretches, *λ*^(*i*)^ (gray bars for the whole assembly, and white bars for *β*_3_). Large strain regions correspond with low values of *λ*^(1)^ and large values of *λ*^(3)^. Note that *λ*^(1)^ *≤* 1, *λ*^(2)^ *≈* 1 and *λ*^(3)^ *≥* 1, which arises from near-incompressibility, i.e. *λ*^(1)^*λ*^(2)^*λ*^(3)^ *≈* 1. **D**. Mapping of stretch *λ*^(1)^ in ATP synthase structure using color scheme in B. Residues of large strain reside in the proton channel, the middle of the peripheral stalk *bb*^*l*^, and the area where *β* and *γ* interact. ATP and ADP appear in green and blue, respectively.

While our PSA framework provides a quantitative and local characterization of rotations, it is unlikely that rotation angle and axis enclose information that is connected to protein function. The key reason is that rotations depend on the spatial alignment of structures, which is arbitrary, and so the value of the local rotation angle does not quantify an intrinsic property of the protein’s deformation. Therefore, we used PSA to quantify strain, which is alignment independent, and thus expected to be closer to a quantification of function itself. Fig. 1C depicts a histogram of the principal stretches, (*λ*^(1)^, *λ*^(2)^, *λ*^(3)^). Since values close to 1 correspond to small deformations, the indication of large strains is to be found in the tails of the histograms. We find that few residues belong to regions that are significantly strained. For instance in *β*_3_ only *∼* 5% of residues are stretched or compressed above 20%, that is *λ*^(1)^ *<* 0.2 or *λ*^(3)^ *>* 1.2.

Figure 1D shows how strains are distributed throughout the structure, with darker red tones denoting higher strain. There are three regions that show significant strain: (*i*) the proton channel at the interface of *a* membrane protein and *c* components of F_o_ (see Fig. 5 for schematics); (*ii*) the central and upper region of the peripheral stalk *bb*′, which has been suggested to be elastically compliant [14, 26, 39, 40]; (*iii*) the central part of *β*_3_, near *γ*, which is where synthesis of ATP takes place. The geometry of high strain regions is very different from that of rotation domains. Whereas rotation domains span large, compact regions, strain accumulates in smaller, disperse functional regions. While some such “strain pockets” are localized at the interface between rotation domains (see below), we emphasize that strain and rotation are not directly correlated, see also Fig. 9 in the SI, as strain characterizes deformations by explicitly excluding rotational information (see Appendix A).

### Within species conservation of *β* strain functional cycle

We focused our analysis on region (*iii*), i.e. *β*_3_ and its interaction with *γ*. Figure 2A shows the distribution of strain found on *β*_3_ and *γ*, as the former undergoes the E *→* D transition with the function of binding ADP. The tip of *γ* shows large strain, due to the large shearing that this small rotating region is subject to. In comparison, a large fraction of strained residues of *β*_3_ are localized at a 2D planar interface between the two rigid domains, identified in Fig. 1A and B, that rotate relative to each other, see Fig. 2B and C. This supports the long-standing hypothesis of strain accumulation in *β* during E *→* D [18]. The involved strained amino acids are found in regions that are distant in space (from the ADP binding pocket G175 to the peripheral K191) and in sequence (such as V440 or A331). Furthermore, we found that the Walker-A motif, a set of eight resides that is evolutionary conserved and functionally critical [41], belongs to region (*iii*) and are clearly over-strained during transitions E *→* D and T *→* E, see Table 1 of the SI. We see, there-fore, that PSA detects mechanically strained regions of proteins that participate in *functionally* important transitions (see section A of the SI for characterization of strain in the *bb*′ stalk).

**TABLE 1:** *Fraction of high strained residues in Walker-A motif of β*. Each row corresponds to a different set of structures (typically different species), and each column to a particular functional transition. In each case the fraction of high strained residues among the eight residues in the Walker-A motif is computed. The average fraction of the chain appears in parenthesis. A strained residue is defined as a residue for which *λ*_1_ *<* 0.2. As one can see, Walker-A motif has an evolutionary conserved pattern of strain, being clearly over and under strained depending on the transition. In particular, we find that in T *→* E the Walker-A motif is very much over-strained.

| Species | E→D | D→T | T→E |
| --- | --- | --- | --- |
| Spinacea oleracea | 0.3 (0.1) | 0.0 (0.0) | 0.8 (0.1) |
| Bacillus sp. PS3 | 0.5 (0.1) | 0.6 (0.1) | 0.1 (0.0) |
| Polytomella sp. pre | 0.3 (0.1) | 0.0 (0.0) | 0.6 (0.1) |
| Ogataea angusta | 0.0 (0.1) | 0.0 (0.0) | 0.5 (0.1) |
| Bos taurus (low-res) | 0.2 (0.2) | 0.2 (0.1) | 0.8 (0.2) |
| Bos taurus (high-res) | 0.3 (0.1) | 0.0 (0.0) | 0.6 (0.1) |
| Mycolicibacterium smegmatis (no drug) | 0.4 (0.1) | 0.0 (0.0) | 0.5 (0.1) |
| Mycolicibacterium smegmatis (drug) | 0.4 (0.1) | 0.0 (0.0) | 0.6 (0.1) |
| Escherichia coli | 0.8 (0.1) | 0.3 (0.1) | 0.1 (0.1) |
| Saccharomyces cerevisiae | 0.6 (0.2) | 0.0 (0.1) | 0.8 (0.2) |

**FIG. 2:**
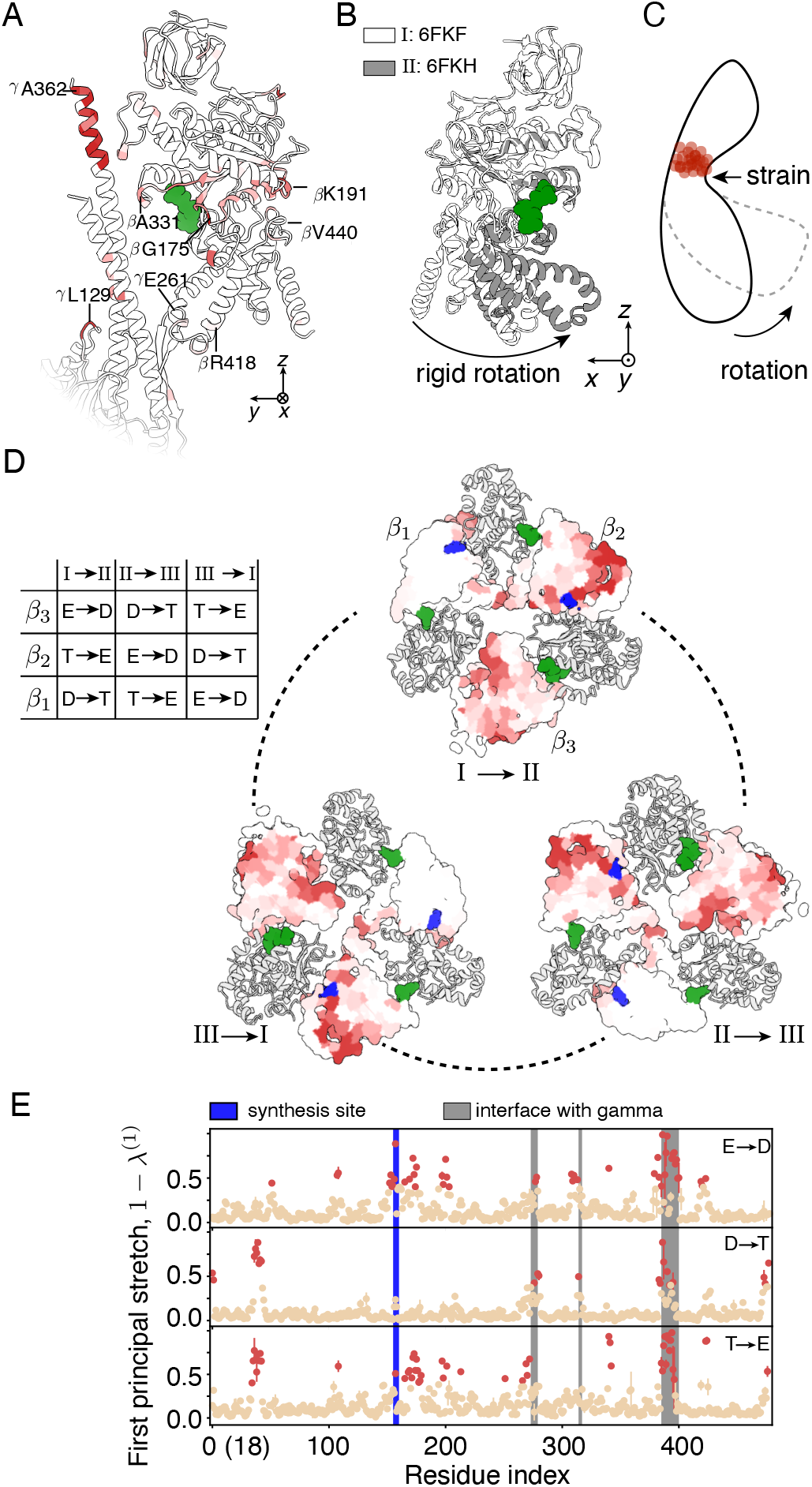
Strain dynamics of β chains through synthesis cycle. **A**. Distribution of strain in *β*_3_ and *γ*. Amino acids in *β* with large strain are located at the interface between the two rotation domains identified in Fig. 1B, near the binding region of ADP. **B**. Superposition of *β* in the reference (I, white) and deformed (II, gray) states. The lower and upper halves of the chain rotate relative to each other as rigid domains. **C**. Schematic representation of the deformation undergone by *β* during E *→* D. **D**. Horizontal cross-section of F_1_ showing strain distribution (color scheme as Fig. 1D, with saturation at *λ*^(1)^ *<* 0.4). High strain accumulates in the plane of ATP synthesis in chains undergoing E *→* D and T *→* E, and in the basal bridge between chains undergoing T *→* E and D *→* T. The table contains the conformational changes undergone by the three *β* chains through a complete cycle of ATP synthesis. **E**. Stretch profiles computed via *λ*^(1)^. Points on each panel depict the mean for the three *β* when they undergo the same transition, with error bars mostly smaller than the point size. Note that profiles characterize each enzymatic transition. The (18) on the *x−*axis denotes labelling of first residue in [14].

In a complete synthesis cycle, each *β* chain performs three distinct functions, related to corresponding structural transition, as summarized by the table in Fig. 2D. While the sequence of the three chains is the same, they are different in their spatial location relative to the peripheral stalk. The stalk in fact breaks the rotational symmetry of F_1_, resulting in a precession of F_1_ relative to F_o_ as the latter rotates. Given these asymmetries, we investigated how well the strain patterns corresponding to each functional transition are conserved across the three *β* chains. In Fig. 2D we show cross-sections of F_1_ at the height of the ADP binding site. Highly strained residues are colored in red, and the rest of them in beige. The strain patterns are characteristic of each transition and are found to be well conserved across the three *β*s. For example, comparing assembly states I *→* II reveals that *β*_2_, which performs the function of ATP release in the transition T *→* E, has a strain pattern that forms a red loop. This red loop is replicated in *β*_1_ and *β*_3_ when they undergo T *→* E. To quantify strain conservation we plotted the strain profiles, i.e. strain values along the sequence, for the three *β*s when they are undergoing each transition, Fig. 2E. The profiles are mostly flat, with peaks localized in small regions. The pattern of peaks is unique for each transitions, acting as the mechanical fingerprint of the corresponding function. For example, there is a peak near the synthesis site when ADP is bound, E *→* D, or ATP released, T *→* E, but in the synthesis step, D *→* T (this may reflect large free-energy changes during nucleotide exchange, compared to moderate ones during bond formation). Furthermore, the strain profiles show a strong degree of conservation across chains for the same structural transition, as reflected by the small standard deviation across the three chains (depicted in the plot as error bars). We conclude that strain in *β* chains is localized in functionally relevant small regions distant from each other, with strain profiles that are characteristic of each enzymatic transition. Furthermore, despite overall precession of F_1_ and other structural asymmetries, these profiles show strong conservation across the three *β* chains when they perform the same function.

### Strain pattern across species

Having established strain as a functionally relevant property, robust to structural asymmetries, we now move to study its evolutionary conservation. To this end we compiled all published structures of ATP synthases, for which a full cycle has been resolved [14, 25–30], and computed the distributions of strain (see section B of the SI for table of supporting files with comprehensive analysis of all structures). Figure 3A shows the spatial distribution of high strain for one third of a cycle on seven different ATP synthases. As one can see these assemblies are structurally very different from one another, which in turn results in distinct distributions of strain. However, in all cases the region (*iii*), in which *γ* interacts with *β*, displays large strain, which parallels the functional conservation of ATP synthase function across species. We thus focused our attention on strain conservation in *β* chains.

**FIG. 3:**
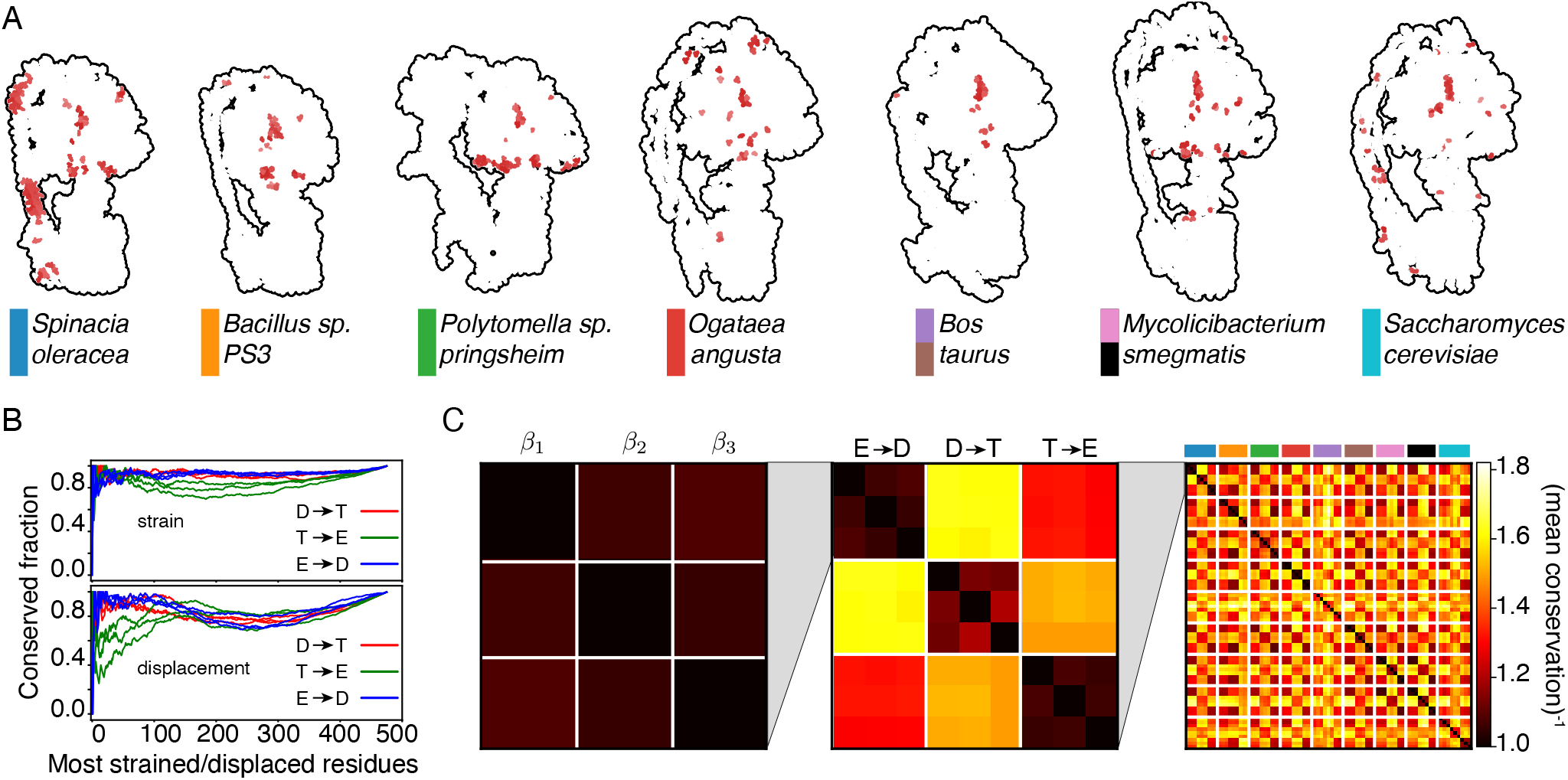
Strain pattern across species. **A**. Strain in ATP synthases of seven different species. Residues sustaining high strain (*λ*^(1)^ *<* 0.25) are depicted in red on top of a silhouette of the corresponding synthase. In all cases high strain accumulates at the height of the *β*, where it interacts with *γ* (region *iii* lies approximately in the same location as in Fig. 1D). Color bars are legends for subsequent panels (note that for *Bos taurus* and *Mycolicibacterium smegmatis* two sets of structures were considered, see section B of the SI for identification of corresponding supporting file). **B**. Fraction of conserved residues between a pair of *β* (of the same species) plotted against the number of residues considered to be strained (obtained by decreasing the threshold separating low-strain from high-strain residues). For all enzymatic transitions, quantifying deformation via strain produces larger conservation than via displacements. **C**. The inverse mean conservation (integral of curves in panel B) can be used to produce a distance matrix that compares different chains (first panel), undergoing different enzymatic transition (second panel), in different species (third panel). Note that darker colors correspond to higher conservation.

As a first step, we confirmed that strain is conserved across *β*s in the same species, spinach chloroplast, when they undergo the same transition, as suggested by Fig. 2D and E. Figure 3B, top panel, shows the fraction of strained residues that are conserved, as a function of the increasing number of residues considered to have high strain. Each curve represents a pair of chains and each color labels a structural transition or function (e.g., a single blue curve corresponds to (*β*_1_, *β*_2_)_E *→*D_, ADP binding): in all cases the conserved fraction is very large. As a comparison to this strain conservation, we also calculated conservation of displacements among the same residues, depicted in the bottom panel. Although they are also significantly conserved, their level of conservation is lower.

Starting from this quantitative measure of strain conservation, we can take the mean conserved fraction (area under each curve in the upper panel of Fig. 3B) as a single number that quantifies how conserved strain is between a pair of chains that undergo a particular enzymatic transition. Figure 3C shows matrices of mean conserved fraction at three different levels: first panel, among the three different chains of the same species that undergo the same transition (this corresponds to Fig. 3B); second panel, among the different chains when they undergo different transitions in the same species; and third panel, among different chains, undergoing different transitions, in different species. Therefore, strain conservation provides a systematic way of comparing different functional transitions of different proteins across different species.

### Evolutionary conservation of strain

Figure 3C quantitatively describes the evolutionary strain conservation in *β* chains participating in ATP synthesis, within seven evolutionarily distant multi-protein assemblies. To elucidate further strain conservation from this multi-dimensional matrix, we used multi-dimensional scaling (MDS) to find its lower dimensional representation. In brief, MDS finds an embedding in low dimensions (two, in this case) such that the distances among points in this embedding are as close as possible to those in the input metric (here the right-most matrix in Fig. 3C).

Figure 4A shows that, when using strain conservation described above as the metric for MDS, enzymatic transitions of different species cluster together: strain patterns that correspond to *the same functional transition* are closer to each other than those that correspond to the same chain, or those of the same species. Importantly, such clustering is *not* observed when using as a metric conservation of residue displacement, Fig. 4B, nor conservation of rotation angle, see Fig. 10 in the SI (in both cases structures were aligned using chain *a* and observables computed using exclusively C_*α*_). Therefore, *strain, and not displacements, is an evolutionarily conserved characteristic of functional transitions in β* chains. It is interesting to note that there is one outlier in the clustering of 4A, corresponding to the ATP synthase from *Bacillus sp. PS3* (orange). Indeed, in [25] it was argued that in this ATP synthase *β* adopts “open-open-close” conformations (as opposed to typical “close-open-close” ones) in order to perform the enzymatic transitions, which naturally results in misclassification of the strain patterns.

**FIG. 4:**
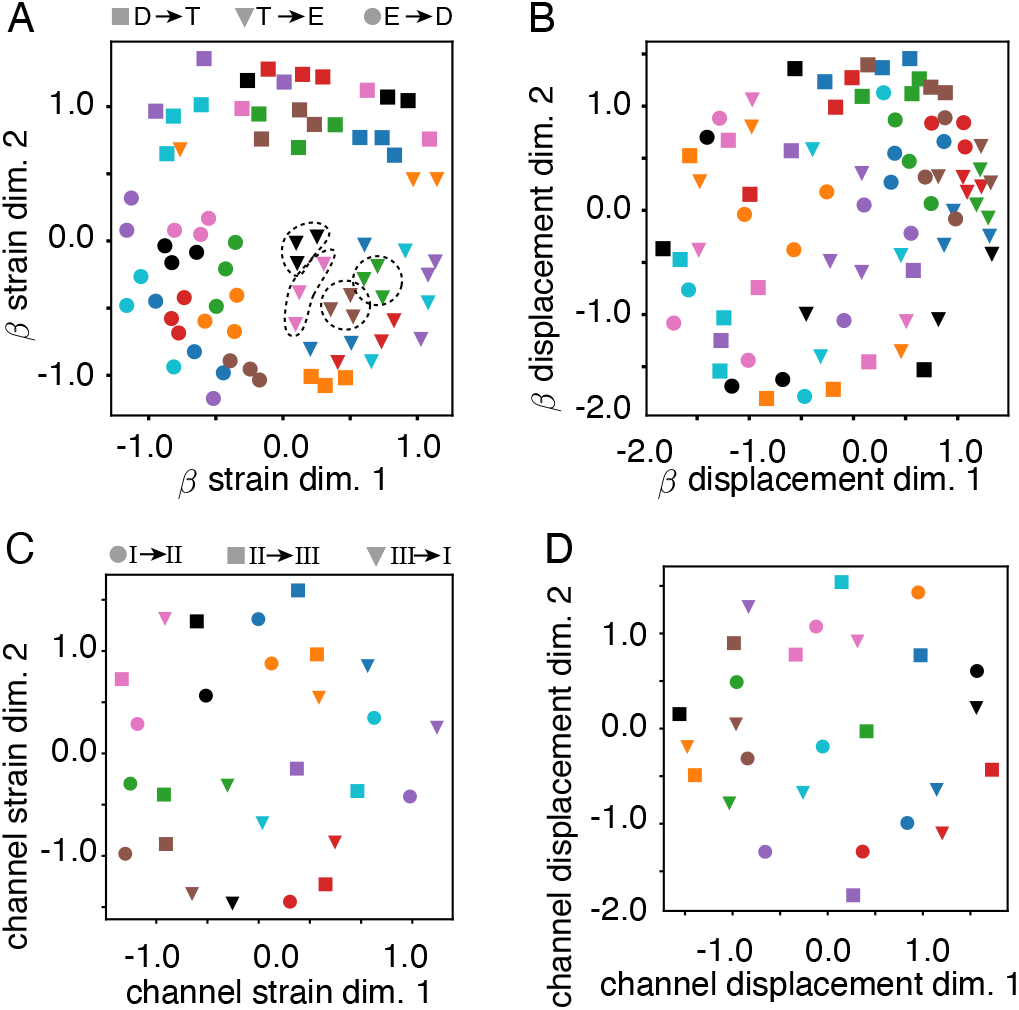
Evolutionary conservation of strain patterns. **A** and **B**. Multidimensional scaling (MDS) of the dissimilarity matrix corresponding to conservation of strain (A) and displacement (B) in *β*. As is readily seen, with few outliers discussed in the text, strain conservation clusters the data according to the enzymatic transitions (gray shades are guides for the eye, and were hand-drawn). This clustering is not observed when displacement conservation is used. Dashed contours highlight examples of within-species conservation, beyond the already mentioned functional conservation. **C** and **D**. Same as A and B but for the proton channel chain *a*. In this case, transitions of each species cluster together when using strain conservation, and they do not when using displacement conservation (as before, gray shades are guides for the eye). Colors correspond to legends in Fig. 3A.

**FIG. 5:**
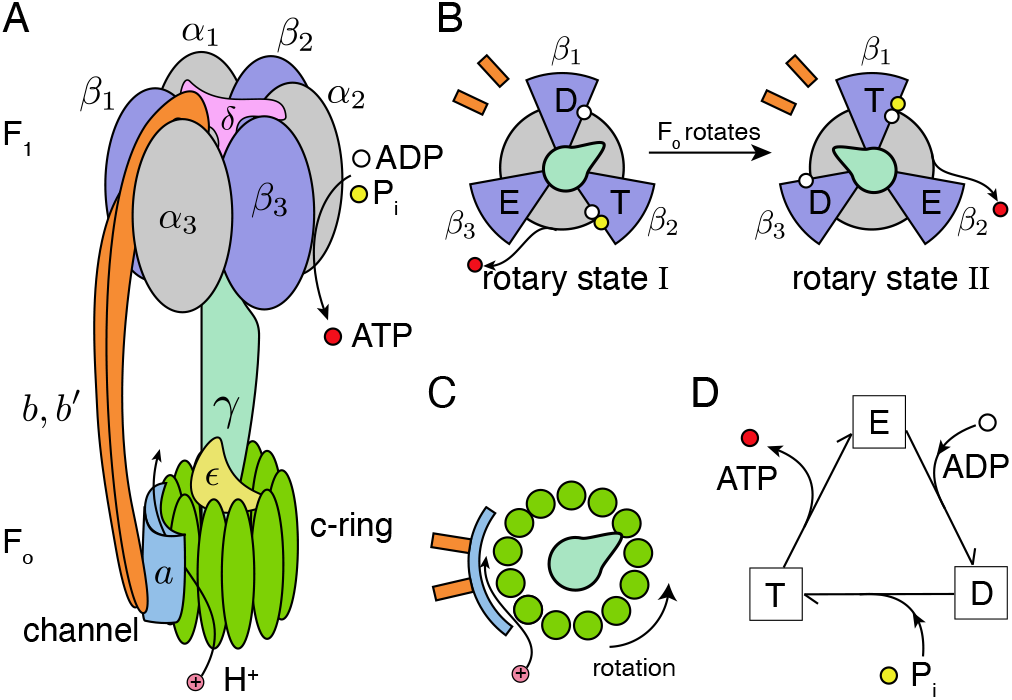
Schematics of the structure and functioning of ATP synthase. **A**. Overview of the ATP synthase protein assembly structure highlighting main components. **B**. Schematic view from the top of the F_1_ complex as F_o_ undergoes a 120° rotations. Each of the three *β*, blue circular sections, adopt a distinct enzymatic state: empty (E), ADP bound (D), and ready for the synthesis of ATP (T). Orange bars represent *bb*^*l*^ and gray circular sections *α*. **C**. Schematic view from the top of the F_o_ complex, indicating rotation direction. **D**. Enzymatic diagram of the synthesis cycle at an individual *β*. Colors of parts are consistent across panels.

**FIG. 6:**
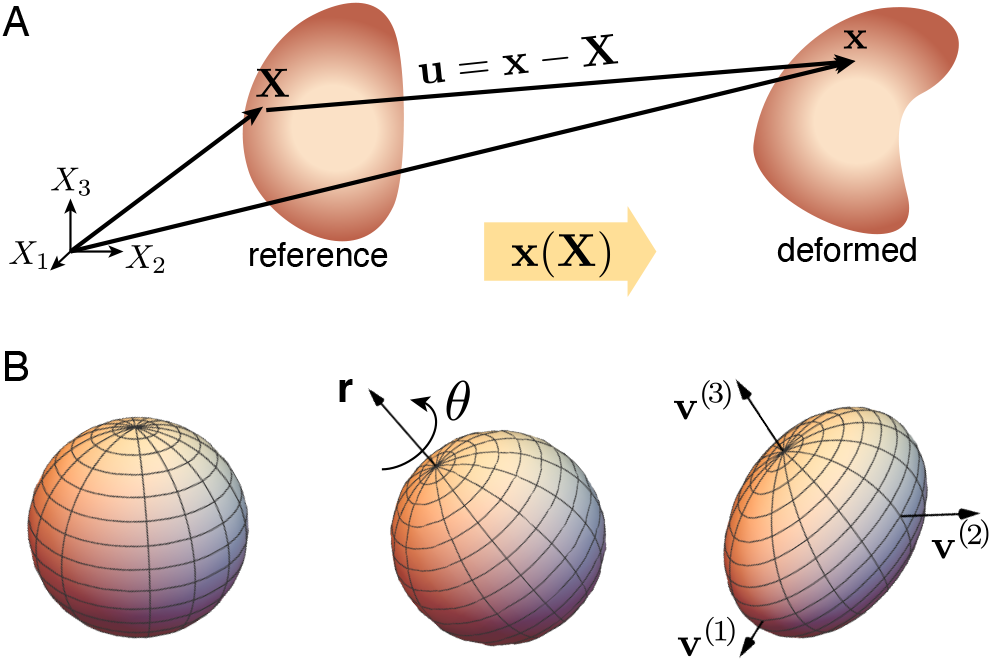
Finite Strain Theory decomposition. **A**. A continuous body with material coordinates **X** in its reference state is described by coordinates **x**(**X**) = **X** + **u** in a deformed state, with **u**(**X**) the displacement field. **B**. The deformation of an elementary sphere inside a material is described by the local rotation angle *θ* around the axis **r** and the principal axes of deformation **v**^(*i*)^ with magnitudes *λ*^(*i*)^ and *i* = 1, 2, 3.

**FIG. 7:**
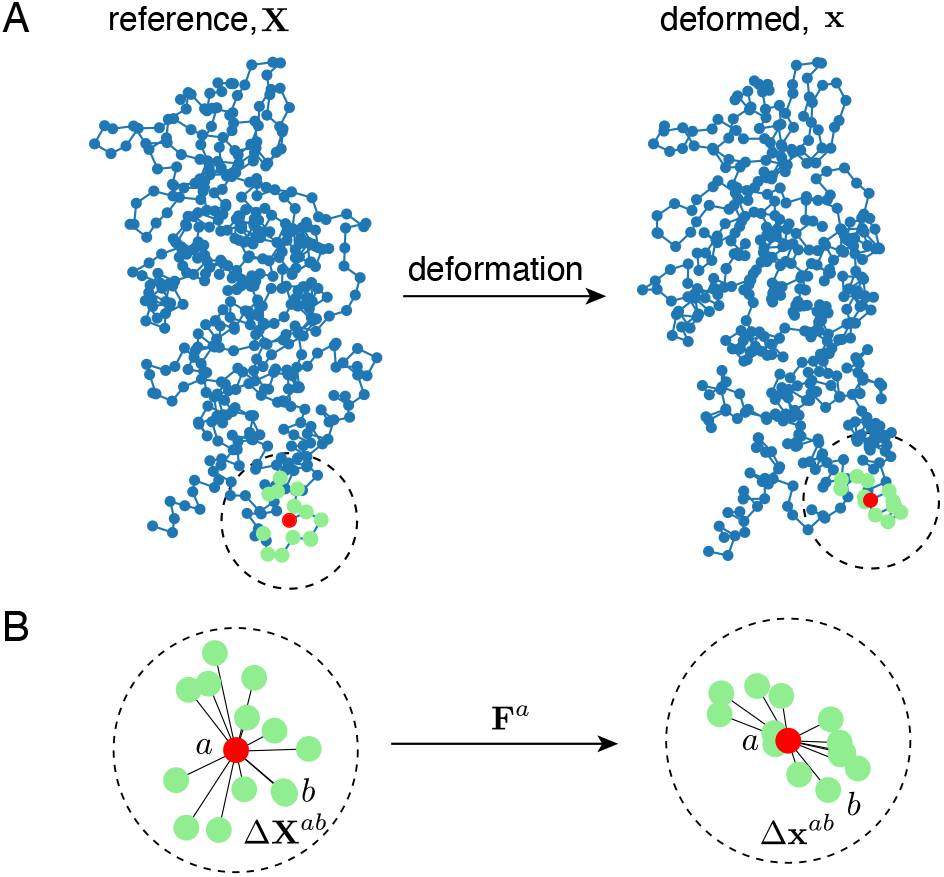
Example of protein deformation. **A**. Two dimensional projection of the C_*α*_ from chain *β* in two conformations, one taken as reference and the other as deformed. A particular atom is highlighted in red, and the surrounding ones within a sphere of radius 8Å in light green. **B**. The atom highlighted in A, labeled *a*, is characterized by distances Δ**X**^*ab*^ to the neighboring *b* atoms in the reference state, and Δ**x**^*ab*^ in the deformed state. The matrix **F**^*a*^ estimates the map of one set of distances to the other, see Eq. D1.

**FIG. 8:**
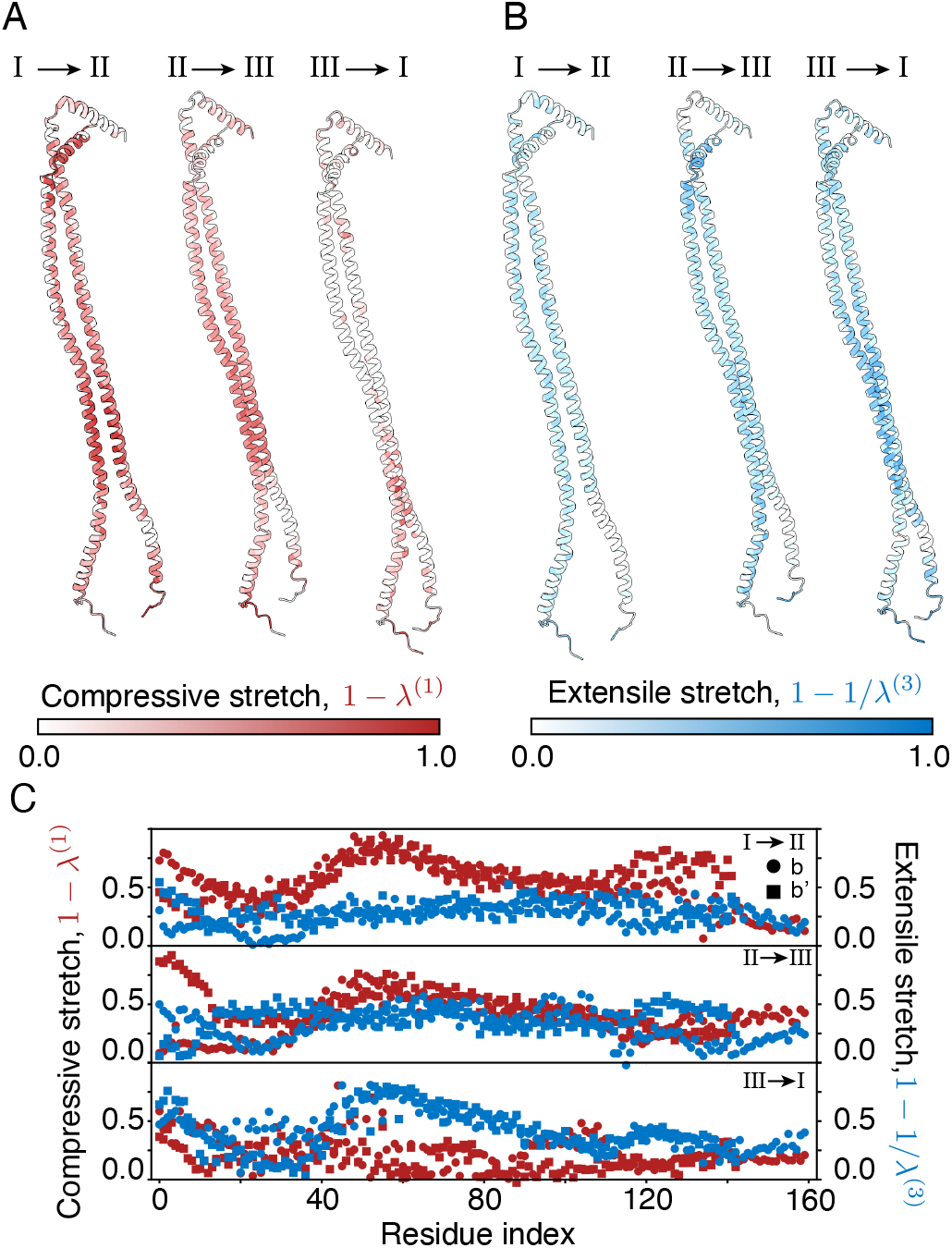
Strain cycle of the peripheral stalk. **A**. Peripheral stalk *bb*^*l*^ colored by compressive strain, where small values of *λ*^(1)^ correspond to strong compressions. As one can see, transition I *→* II implies a strong compression, transition II *→* III intermediate, and the transition III *→* I shows almost no compression. **B**. Same as A but with the extensile strain. The results mirror those of A, with extensile strain being contained in III *→* I, with the other two transitions containing small extensile strain. **C**. Compressive and extensile strain profiles of the two chains in the peripheral stalk for all three transitions. Whereas in II *→* III extension and compression domains are similar, in I *→* II and III *→* I either one or the other dominates. Both chains, *b* and *b*^*l*^, exhibit similar behavior.

**FIG. 9:**
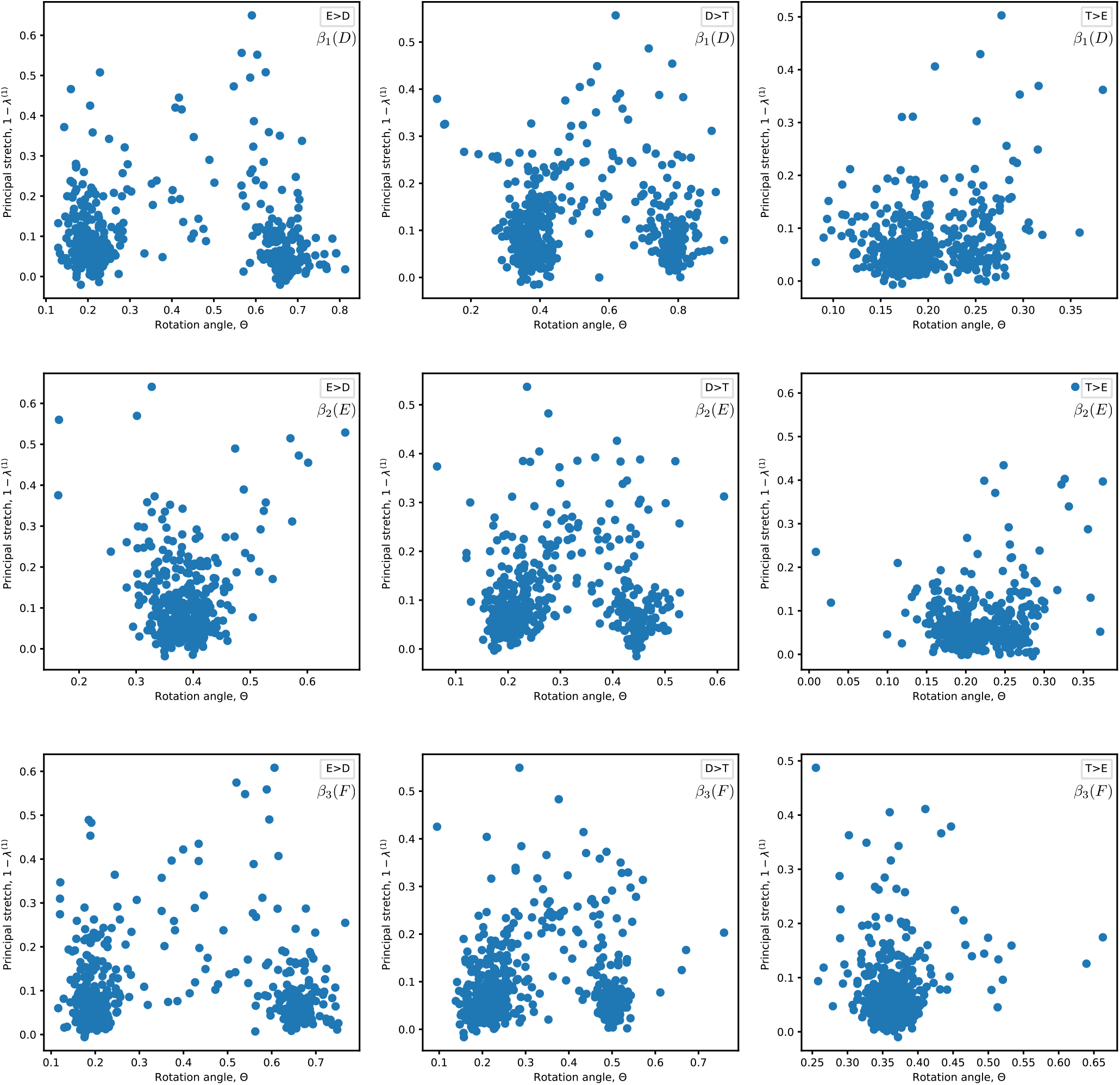
Relationship between strain and rotation for β. Scatter plot of the principal stretches against the rotation angle for all residues in the case of *Bacillus sp. PS3* in each of the three functional transition (columns) for each of the three *β* chains (rows). No correlation is apparent. Similar results were also obtained for other species.

**FIG. 10:**
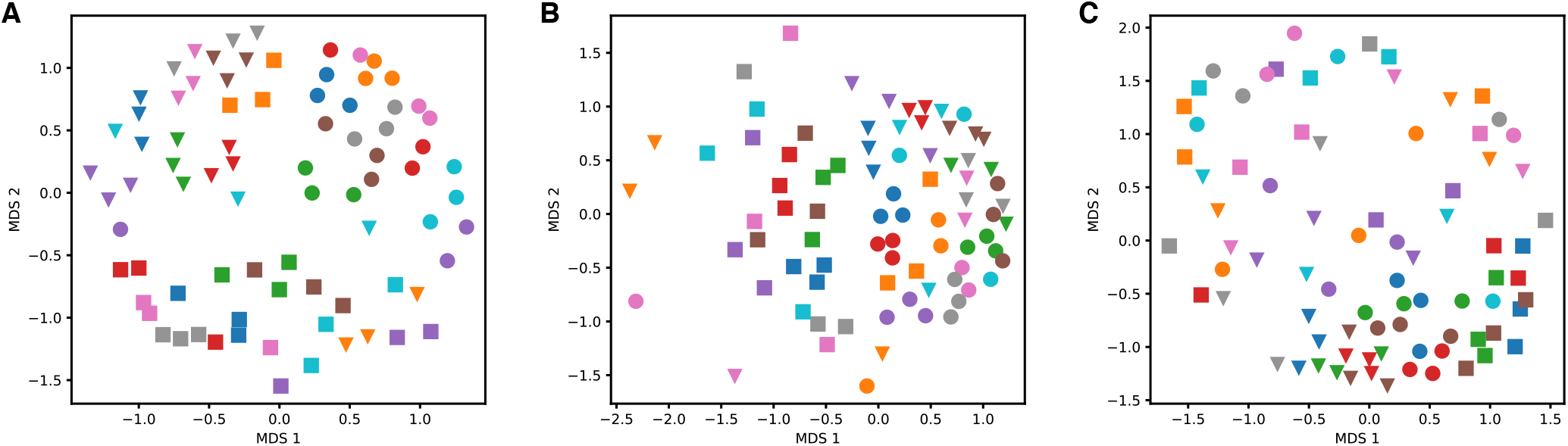
Comparison of conservation analysis on β chain using strain, rotation angles, and displacements. **A**. Multi-dimensional-scaling projection in two dimensions of the conservation dissimilarity matrix described in the main text. This is the same as Fig. 4A, and exhibits clustering of functional transitions. **B**. Same as as A, but in this case the conservation matrix was used using rotation angle profiles instead of strain profiles. That is, instead of profiles such as those in Fig. 2E, analogous ones were used in which the vertical axis displays the local rotation angle. Such profiles were then used as the basis for the conservation analysis. In this case, we do not observe clear clustering of functional transitions. **C**. Same as A and B, but displacement profiles were used to construct the conservation matrix. That is, the distance between the position of each residue in one conformation to the position of that same residue in the other. As in B, clustering is not observed. In all cases only C_*α*_ were used, and structures were aligned using chain *a*. Legends are the same as in main text.

Remarkably inside each functional cluster, we also observe a large degree of within species clustering, see contours shapes in Fig 4A. As for the functional clustering, for this within-species clustering there seem to be some notable exceptions. Most notably, the *Bos taurus* dataset (in violet) is very spread out. It is likely that this may arise from a low resolution of the corresponding cryo-EM structure [28]. This second level of clustering indicates that species-specific adaptations are indeed present for each component of the cycle but become only apparent once the dominant functional/structural effect has been first taken into account.

Our analysis of strain conservation can be extended to other relevant regions of the ATP synthase. For instance, an analogous MDS embedding performed on the proton channel protein *a* shows that all transitions of the same species can be better clustered together while using strain conservation, Fig. 4C, than by using displacement conservation as a metric, Fig. 4D. Of the two outliers seen in Fig. 4C, one corresponds again to low resolution *Bos taurus* structures. The other outlier, depicted in black, corresponds to the structures of *Mycolicibac-terium smegmatis*, obtained in presence of the inhibitory drug bedaquiline (BDQ) [31]. BDQ binds to the c-ring and is known to affect the interaction between the c-ring and the channel protein *a*. In agreement with this, we found that pockets of strain characterizing the deformations of the *a* protein in absence of BDQ, were suppressed in presence of BDQ, see Fig. 11 in the SI. Once again, as in the case of *β* chains in *Bacillus sp. PS3*, our strain-based quantitative analysis captures subtle differences in structural transitions in an unbiased manner. Overall, the analysis of the proton channel further supports that species-specific adaptations are revealed when using a function-related metric, as we have shown strain is.

**FIG. 11:**
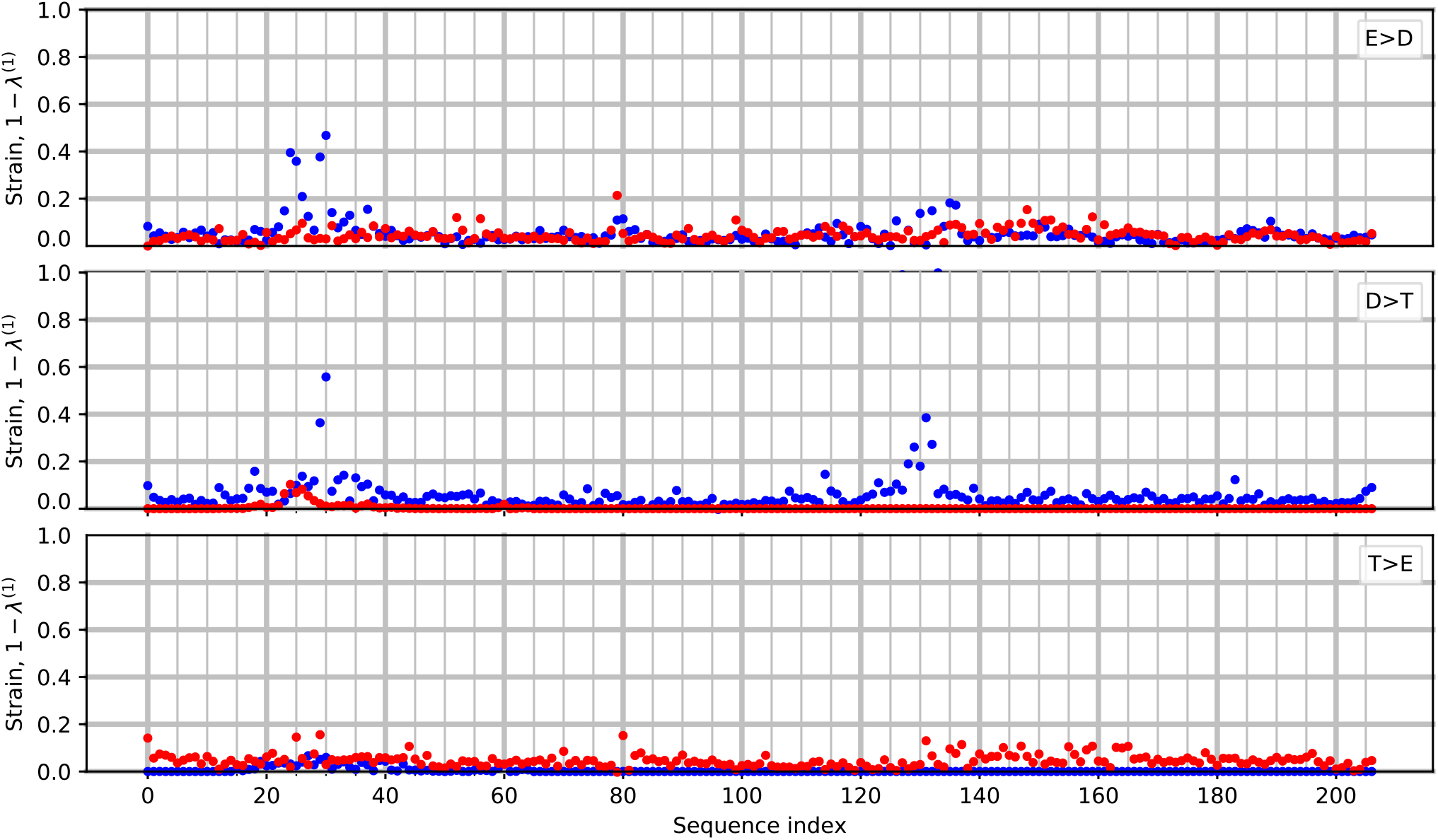
Strain pattern of channel protein in response to drug treatment. The strain pattern of the channel protein *a* of *M. smegmatis* in presence (red) and absence (blue) of the inhibiting drug BDQ is shown for each of the three functional transitions. As one can see, in absence of BDQ three different patterns distinguish the functional transitions. Once function of the ATP synthase is abolished by BDQ the patterns become suppressed.

In conclusion, strain profiles can be used for functional evolutionary analysis as they quantify conservation properties (as well as point to subtle differences) better than measures based on residues displacements.

## Discussion

Evolution of proteins is associated with changes in their encoded sequence, translating directly into changes of their three-dimensional structure. In turn, these changes in structure, together with the changes taking place for all cellular partners, modify the structural changes that a protein can undergo. The set of the allowed functional changes in structure can be viewed as the protein’s “molecular phenotype”, which encompasses for instance its allosteric or enzymatic qualities. Therefore, quantification of functional transitions in protein structures is crucial to provide a quantitative assessment of protein genotype-phenotype maps.

The recent explosion of cryo-EM data has resulted in a surge of methods aimed at interpreting ensembles of electron microscopy images [42, 43] and reconstructing protein atomic maps [44–46]. In contrast, quantitative methods for the comparative analysis of such maps are still underdeveloped. Most existing approaches come from the molecular dynamics literature [47, 48], and require large sets of simulation-generated atomic maps to perform low-dimensional embeddings. This requirement renders them inadequate for the handful atomic maps that are experimentally accessible. Instead, the closest approach to protein strain analysis (PSA) is normal mode analysis (NMA) [49, 50], which characterizes a structure by the normal modes of its elastic network model. However, while NMA characterizes fluctuations of a single structure by displacement fields, PSA characterizes potentially large structural transitions between a pair of structures by strain fields, which are alignment independent. Therefore, PSA presents a new paradigm to compare pairs of atomic maps grounded on fundamental physical principles.

We have shown that strain shows a large extent of evolutionary conservation for the same function. This implies that mechanical strain is not only an important functional, but also *evolutionary* quantity. Because our analysis corresponds to structural changes of the same genotype (for each species), the concept of conservation we have discussed so far is different from that of sequence conservation. Nevertheless, given the relevance of sequence conservation in determining the structure of proteins [51], an important question is how strain relates to sequence conservation. To address this we performed multiple sequence alignment on the set of all *β* chains studied in our work, and quantified conservation by the Shannon entropy on the position-specific-score-matrix, see Fig. 12 in the SI. Remarkably, we found no correlation between the Shannon entropy and the magnitude of strain. This suggests that strain patterns are not a direct consequence of sequence conservation, and instead reflect a convergent phenotypic trait in the light of genotypic variations.

**FIG. 12:**
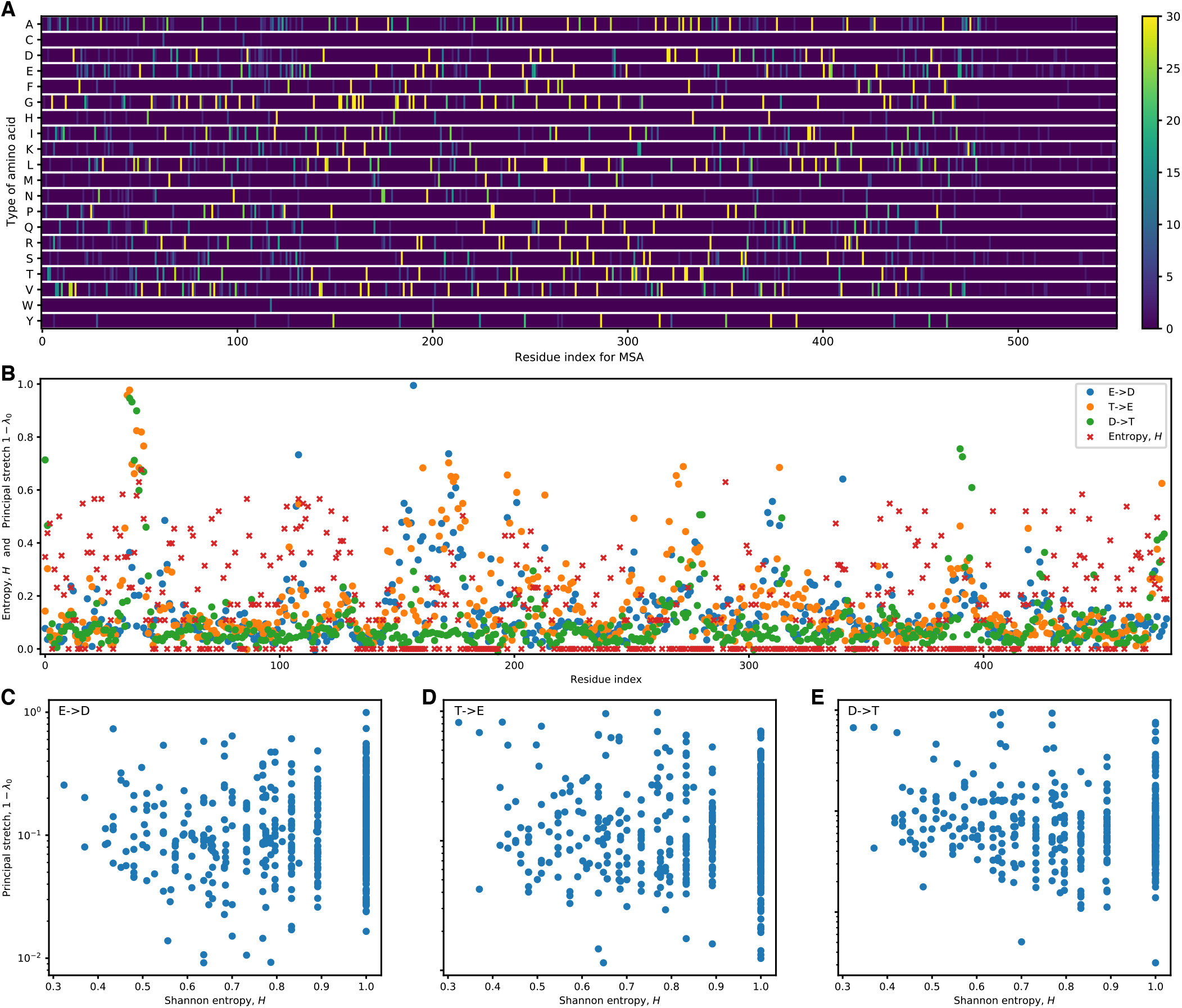
Relation between residue conservation and high strain in β chain of Spinach cholorplasts. **A**. Position-specific-score-matrix (PSSM) after performing MSA in all chains of the main text. Color bar represents number of chains in which residue is conserved. **B**. Shannon entropy for each residue, which measures conservation, as well as the first principal stretch in each of the three transitions, which measures strain. **C, D, E**. Scatter plot of Shannon entropy against the first principal stretch for transitions E *→*D, T *→*E, and D *→*T, respectively. No correlation is apparent.

In conclusion, strain emerges as an evolutionarily conserved quantity that characterizes well the nature of functional transitions in the ATP synthase. This should not be surprising: after all, proteins are *(visco-)elastic evolved materials* so it is natural that strain, introduced by elasticity theory, should be used to better understand their functioning and their evolution.

## Appendix A Protein Strain Analysis in a nutshell

We now summarize the key aspects of finite strain theory (FST) [9, 15], and then elaborate on how to adapt FST for the analysis of proteins, which we refer to as protein strain analysis (PSA). The main aim of the FST formalism, summarized in C, is constructing the (Lagrangian) strain tensor *γ*_*ij*_ at every point of a material. This mathematical object quantifies the difference between measuring distances in a reference and a deformed structure. In other words, 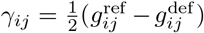, where the metric *g*_*ij*_ quantifies how distances in a material measured in the direction of coordinate *i* change along coordinate *j* [52]. As distances are coordinate-independent properties, so is the strain tensor. In addition, FST also provides information on the local rotation angle *θ* and rotation axis **r**, which are however alignment dependent. We remark that the strain tensor ***γ*** does not depend on an elasticity model. In contrast, to discuss stress a strain-stress relation needs to be defined. For hyper-elastic solids this can be done from a particular elastic energy density (e.g. Saint-Venant, neo-Hookean, etc.).

In our work, we characterize the strain tensor by the principal axes of deformation **v**^(*i*)^ (with *i* = 1, 2, 3). The corresponding principal stretches along the three axes, *λ*^(*i*)^, which are, importantly, invariant under coordinate transformations, quantify the amount of deformation (note that *λ*^(*i*)^ = 1 corresponds to lack of deformation). Intuitively, a local deformation can be thought of as transforming a small sphere of material into a small ellipsoid: **v**^(*i*)^ correspond then to the direction the ellipsoid axes and *λ*^(*i*)^ to their relative stretch (Fig. 6). We emphasize that strain is a rate of change in distances, and not rotations. While rotations in a material can indirectly and non-locally affect strain, there is no a priori direct correlation between the two (see Fig. 9 in the SI).

In Protein Strain Analysis, PSA, we apply FST to protein structural changes. To this end, we first identify the positions of all the carbon atoms C_*α*_ in the reference state, **X**, and deformed state, **x**. This information is obtained from atomic maps, readily available in the form of `pdb` or `cif` files. The next step is to calculate the deformation gradient **F**, which involves derivatives of **x** with respect to **X**. Because we are attempting to compute, **F**, which is a continuum concept, in a discrete material, an approximation method is needed. Different such approaches have been developed in the literature of discrete materials [53, 54] (see D for an overview of three such methods). In this paper, we use the method of Ref. [53] that estimates **F** at a given atomic position by considering all neighboring atoms within a sphere of radius *r*. Once the deformation gradient field is calculated, deducing the rotation and strain features can be done as described in C. In order to facilitate the usage of PSA by the research community, we have created a `Python 3` library that contains a detailed implementation of PSA, https://github.com/Sartori-Lab/PSA.

## Appendix B: Schematics of the ATP synthase structure

The ATP synthase protein assembly consists of two motor complexes, F_o_ and F_1_, that oppose each other, see Fig. 5. The F_o_ complex is constituted by the *c −* ring (immersed in the membrane) and the central “stalk” *γ* and *E*. F_o_ rotates driven by a controlled flux of protons through the channel between the *c*-ring and *a*, which acts as a stator. The F_1_ complex is prevented from rotating by the stalk *bb*′ with which it interacts via *δ*. As a consequence, rotation of *γ* triggers sequential conformational changes of the three *β*, which participate in the synthesis of ATP from ADP and inorganic phosphate P_i_.

One important structural feature of the ATP synthase assembly is that the number of protein monomers in the c-ring is not always a multiple of 3, and so there is a stoichiometric mismatch between F_o_ and F_1_. In addition, the rotation of gamma is not homogeneous, but occurs through three discrete jumps. These three jumps do not span exactly 120° each, likely due to the aforementioned mismatch in stoichiometry.

## Appendix C: Summary of Finite Strain Theory

A body characterized by material cartesian coordinates *X*_*i*_, with *i* = 1, 2, 3, is deformed to **x**(**X**), see Fig. 6A. The *deformation gradient F*_*ij*_(**X**) is then given by

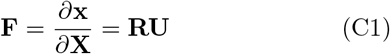

where in the second equation we used the polar decomposition to obtain the *rotation matrix R*_*ij*_(**X**) and the *right stretch tensor U*_*ij*_(**X**). The definition of non-linear *Lagrangian strain tensor γ*_*ij*_(**X**) follows directly:

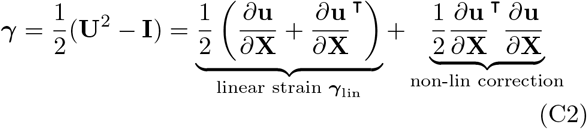

where we used the *displacement field u*_*i*_ = *x*_*i*_ *− X*_*i*_, see schematic below.

The local *rotation angle θ*(**X**) and *axis r*_*i*_(**X**) are calculated from the rotation matrix via

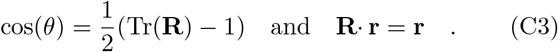

The *principal stretches λ*^(*i*)^(**X**) and *principal axes* **v**^(*i*)^(**X**) of deformation are obtained from the eigen-problem

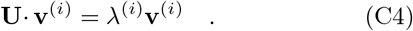

The *λ*^(*i*)^, custimarily ordered by increasing magnitude, are related to the eigenvalues of the Lagrangian strain tensor through *γ*^(*i*)^ = ((*λ*^(*i*)^)^2^ *−* 1)*/*2.

Overall, Finite Strain Theory allows to decompose a deformation **x**(**X**) into a global translation, local rotations (characterized by **r** and *θ*) and local strain (characterized by **v**^(*i*)^ and *λ*^(*i*)^), see Fig. 6B. We remark that **F, R** and **U** are invariant under rigid body translations of either reference or deformed structures. **U** and ***γ*** are additionally invariant under finite angle rotations, and *λ*^(*i*)^ are, furthermore, coordinate invariant.

## Appendix D: Deformation gradients in discrete materials

Consider a protein that transitions from a reference state (with atomic positions **X**) to a deformed state (with atomic positions **x**), for which we want to perform FST. To estimate at a given atom *a* the deformation gradient, **F**^*a*^, we consider all atoms, *b*, within a sphere of radius *r* centered at *a*, see Fig. 7.

For the reference conformation we define vectors connecting the target atom *a* with its neighbors *b*, Δ**x**_*ab*_, and analogously for the deformed state, Δ**X**^*ab*^. A second order approximation of the deformation gradient satisfies

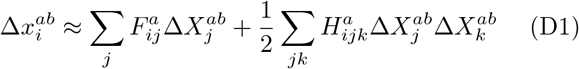

for each (*ab*) pair.

Different algorithms, which we now summarize, attempt at estimating **F**_*a*_ from the expression above.

### Minimal

**H**^*a*^ is set to zero, which keeps the approximation linear, see [53]. Furthermore, only the three *b* neighbors closest to *a* are considered. Since on the previous equation each neighbor adds three conditions on the components of **F**^*a*^, which has nine components, this is sufficient to determine a unique **F**^*a*^.

### First order

Still **H**^*a*^ is set to zero, but all *b* neighbors are kept, which leads to an over-determined system for **F**^*a*^ when there are more than 3 neighbors. An error function is defined *ϕ*(**F**^*a*^) = Σ_*b*_ || Δ**x**^*ab*^ *−* **F**^*a*^ Δ**X**^*ab*^|| ^2^. The minimum of this function is taken as an estimator for **F**^*a*^, see [53].

### Second order

Analogous to the previous, but the error function takes into account the second order corrections. The system is overdetermined for more than 12 neighbors, and the computational cost is substantially larger. The gain is a discrete strain tensor that satisfies compatibility conditions [54].

## Acknowledgments

*Acknowledgements*. The authors would like to thank W. Kühlbrandt, H. Guo, J.L. Rubinstein, J. Howard, and T. Tlusty for helpful discussions, as well as E. Kussell and V.H. Mello for a critical reading of the manuscript. The authors would also like to thank V.H. Mello for support in the design of Box 2 in the supplementary material. This work was partly funded by a laCaixa grant to PS. Early stages of this research have been partly supported by grants from the Simons Foundation to S.L. through the Rockefeller University (Grant 345430) and the Institute for Advanced Study (Grant 345801).

## V. SUPPLEMENTARY MATERIAL

### A. Extension-compression cycle in peripheral stalk

So far we have discussed strain on the catalytic *β*. We now focus instead on the peripheral stalk, which has been suggested to be responsible for energy storage via elastic deformation. Figure 1D revealed a large accumulation of strain on *bb*′ during assembly transition I *→*II, see also Fig. 8A. This corresponds to compressive strain, i.e. to a region with low values of *λ*^(1)^. In contrast, Fig. 8B shows that the same I *→*II transition does not contain significant extensile strain, large values of *λ*^(3)^. The converse is true for the transition III *→*I, which corresponds to significant extensile strain, but very little compressive strain. Finally, during the II *→*III transition both extensile and compressive strain are low. Therefore, throughout a full cycle, the peripheral stalk undergoes compression (I *→*II), bending (II *→*III), and extension (III *→*I) transitions.

Figure. 8C allows to quantify this cycle. Indeed, we observe that compressive strain dominates in I *→* II, whereas extensile strain dominates in III *→* I. It is worth noting that both chains, *b* and *b*′, behave similarly, which speaks to the fact that strain is a local, yet coarse-grained, characterization of deformations. Taken together, our analysis shows that the peripheral stalk undergoes a deformation cycle that may allow for storage and release of energy during ATP synthesis. Such elastic energy accumulation has been suggested to mediate the loose coupling between F_o_ and F_1_ [14, 39, 40], which allows to accommodate their stoichiometric mismatch: 3*αβ* compared to14 chains in c-ring.

### B. Description of additional materials

We append all the `jupyter` notebooks necessary to reproduce the results in the main text, see table below. The notebooks make use of `PyPSA.` Except `Strain evolution.ipynb`, all other notebooks correspond to a detailed analysis of a pair of `pdb` files that were contributed in the corresponding paper. Since all papers contain, at least, three structures, at the beginning of the notebook there is an option to choose among the corresponding `pdb` files. There are also options to highlight diverse chains (chain *β*_3_ is the one highlighted by default). Therefore, while in the main text only a detailed analysis of the spinach chloroplast structures in [14] is presented, the notebooks contain analogous analysis for the structures of all the other species.

The notebook `strain evolution.ipynb` contains data structures that standardize the nomenclature of chains, assembly states, and enzymatic transitions. This is necessary in order to perform comparative analysis across species. Finally, the notebook `strain evolution.ipynb` corresponds to the evolutionary analysis behind Figures 3 and fig:straincons. We note that this second notebook loads the output of `strain evolution.ipynb`.

| File name | Description |
| --- | --- |
| `Hahn_2018.ipynb` | Detailed analysis of structures in [14] |
| `Guo_2019.ipynb` | Detailed analysis of structures in [25] |
| `Guo_2021.ipynb` | Detailed analysis of structures in [31] |
| `Guo_2022.ipynb` | Detailed analysis of structures in [32] |
| `Murphy_2019.ipynb` | Detailed analysis of structures in [26] |
| `Sobti_2020.ipynb` | Detailed analysis of structures in [30] |
| `Spikes_2020.ipynb` | Detailed analysis of structures in [29] |
| `Vinothkumar_2016.ipynb` | Detailed analysis of structures in [27] |
| `Zhou_2016.ipynb` | Detailed analysis of structures in [28] |
| `reference_tables.ipynb` | Standardizes the labels for chains and assembly states |
| `strain_evolution.ipynb` | Evolutionary analysis of data presented in all papers above |
| `analyze_msa.ipynb` | Analysis that compares strain to sequence conservation |

### C. Additional figures

